# Neutralising antibodies against the SARS-CoV-2 Delta variant induced by Alhydroxyquim-II-adjuvanted trimeric spike antigens

**DOI:** 10.1101/2021.08.18.456891

**Authors:** Claudio Counoupas, Paco Pino, Alberto O. Stella, Caroline Ashley, Hannah Lukeman, Nayan D. Bhattacharyya, Takuya Tada, Stephanie Anchisi, Charles Metayer, Jacopo Martinis, Anupriya Aggarwal, Belinda M. Dcosta, Joeri Kint, Maria J Wurm, Nathaniel R. Landau, Megan Steain, Stuart G Turville, Florian M Wurm, Sunil A. David, James A. Triccas

**Affiliations:** School of Medical Sciences, Faculty of Medicine and Health, The University of Sydney, Camperdown, NSW, Australia; Tuberculosis Research Program, Centenary Institute, Sydney, NSW, Australia; ExcellGene SA, 1870 Monthey, Switzerland; Kirby Institute, University of New South Wales, Sydney, NSW, Australia; Department of Microbiology, NYU Grossman School of Medicine, New York, New York, USA; Life Science Faculty, Swiss Federal Institute of Technology Lausanne (EPFL), 1015 Lausanne, Switzerland; Virovax, Lawrence, Kansas 66047, United States; Sydney Institute for Infectious Diseases and the Charles Perkins Centre, The University of Sydney, Camperdown, NSW, Australia

## Abstract

Global control of COVID-19 will require the deployment of vaccines capable of inducing long-term protective immunity against SARS-CoV-2 variants. In this report, we describe an adjuvanted subunit candidate vaccine that affords elevated, sustained and cross-variant SARS-CoV-2 neutralising antibodies (NAbs) in multiple animal models. Alhydroxiquim-II is a TLR7/8 small-molecule agonist chemisorbed on aluminium hydroxide. Vaccination with Alhydroxiquim-II combined with a stabilized, trimeric form of the SARS-CoV-2 spike protein (termed CoVac-II) resulted in high-titre NAbs in mice, with no decay in responses over an 8-month period. NAbs from sera of CoVac-II-immunized mice, horses and rabbits were broadly neutralising against SARS-CoV-2 variants. Boosting long-term CoVac-II-immunized mice with adjuvanted spike protein from the Beta variant markedly increased levels of NAb titres against multiple SARS-CoV-2 variants; notably high titres against the Delta variant were observed. These data strongly support the clinical assessment of Alhydroxiquim-II-adjuvanted spike proteins to protect against SARS-CoV-2 variants of concern.

## MAIN TEXT

COVID-19 vaccines have had a remarkable impact on controlling the pandemic in high and middle-income countries. However, global access to affordable COVID-19 vaccines remains a critical issue^1^. Neutralising antibodies (NAbs) are considered the key determinant of SARS-CoV-2 protective immunity^2,3^, yet in both natural infection and vaccination the levels of NAbs decay over time^4,5^. This issue is compounded by the emergence of SARS-CoV-2 variants circulating that show partial resistance to current vaccines^6,7^, highlighting the need for next-generation vaccines that display strong and persistent immunity.

Subunit vaccines represent a safe, affordable platform for COVID-19 vaccines, with positive outcomes for some candidates from clinical trials^8,9^. To develop a subunit vaccine that is suitable for global distribution, large-scale antigen manufacture and selection of adjuvants are critical issues. To address these issues, we used stabilized, trimeric SARS-CoV-2 spike protein produced with a fully scalable, chemically defined and GMP-compliant production process^10^. To create the CoVac-II vaccine candidate, trimeric spike protein was combined with Alhydroxiquim-II (AHQ-II); a small molecule toll-like receptor 7/8 agonist that is chemisorbed to alum. AHQ-II is the adjuvant used in the Covaxin COVID-19 vaccine, which has received emergency use approval in multiple countries^11^. Importantly, recombinant spike trimer expressed and purified from Chinese hamster ovary (CHO) cells was stable when stored at multiple temperatures and resistant to repeated freeze-thaw cycles (Fig. S1). Following vaccination of mice with CoVac-II (Fig. 1A), high-titre NAbs were apparent in plasma as early as 2 weeks after the first immunization (Fig. 1B). NAb titres did not drop over the time course and reached their peak at 252 days post-vaccination. This corresponds with the excellent stability of both the spike antigen (Fig. S1) and adjuvant used; AHQ-II is stable for at least one year at room temperature (not shown). When compared to spike alone, AHQ-II increased Nab titres by approximately 1000-fold; this increase compares favourably to the effect seen with adjuvants Matrix-M (approximately 10-fold increase) and AS03 (approximately 500-fold) used in other SARS-CoV-2 subunit vaccines^12,13^. Spike plus Alhydrogel (SpK^Alum^) resulted in NAb titres well above that elicited by spike alone, however titres were approximately 100-fold lower at day 252 compared to those from CoVac-II-vaccinated animals (Fig.1B).

**Figure 1.**
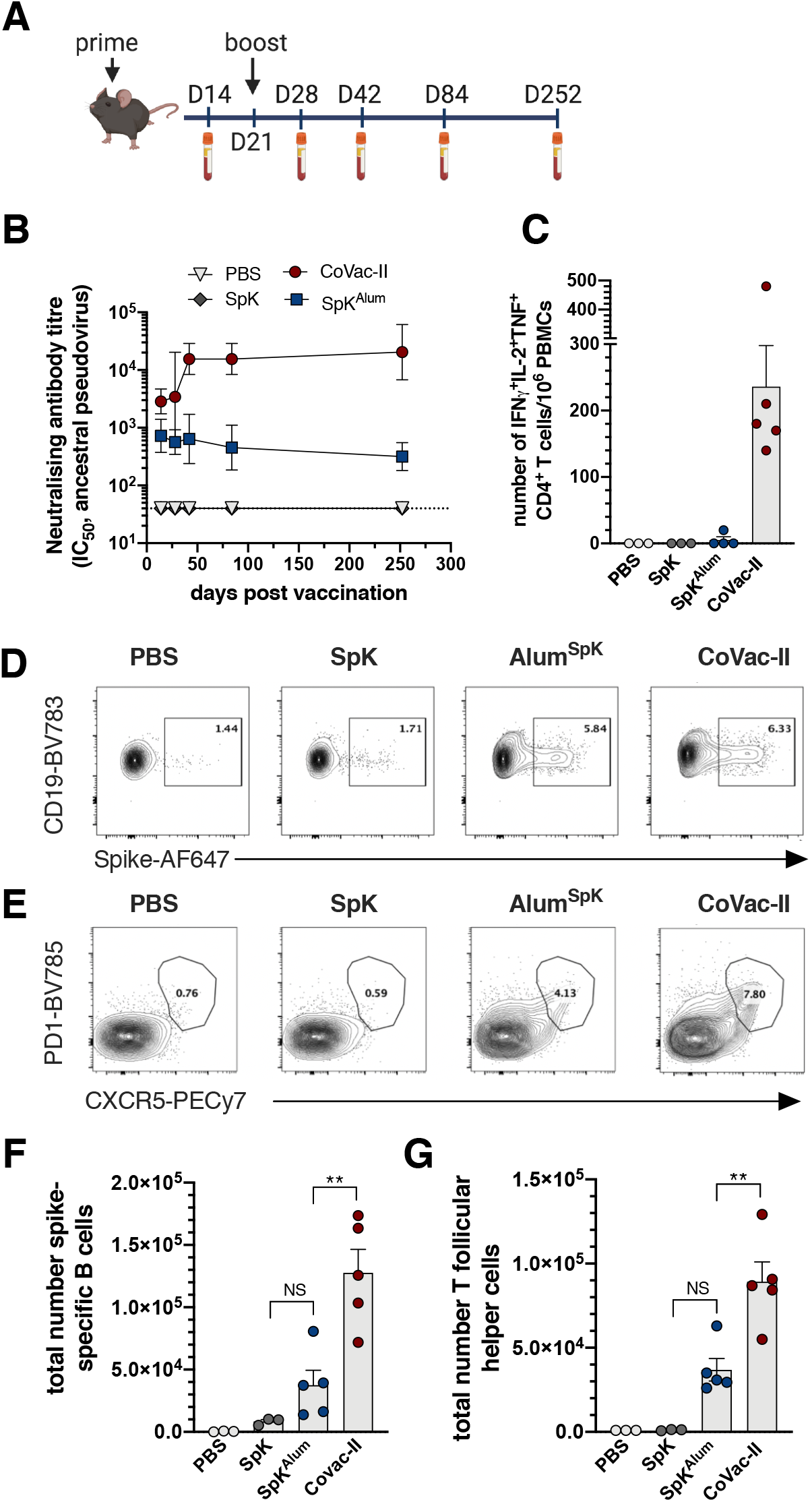
Sustained neutralising antibody titres and generation of multifunctional CD4^+^ T cells responses after vaccination with Alhydroxiquim-II-adjuvanted spike antigen. **A**. C57BL/6 mice were vaccinated s.c at day 1 and day 21 with PBS, SpK (5 μg ancestral spike), SpK^Alum^ (5 μg spike/100 μg Alhydrogel) or CoVac-II (5 μg spike/100 μg Alhydroxiquim-II). **B.** Neutralising antibody (NAb) titres (IC_50_) in plasma were determined using ancestral spike-pseudotyped virus. Dotted line shows the limit of detection. **C.** PBMCs taken 1 week post-boost were restimulated with 5 μg/mL of SARS-CoV-2 spike and the number of cytokine-expressing CD4^+^ T cells determined by flow cytometry. **D-G**. C57BL/6 mice were vaccinated as in **A** and 7 days the frequency of spike-specific B cells (**D**, CD19^+^MHCII^+^) or T follicular helper T cells (Tfh) (**E**, CXCR5^+^PD-1^+^) determined by flow cytometry. The total number of spike^+^ B cells (**F**) and Tfh cells (**G**) is also shown. Data presented as geometric mean ± geometric SD (**B**) or mean ± SEM (C, F, G). Significant differences between groups were determined by one-way ANOVA.

To determine the pattern of T cell immunity induced by the CoVac-II vaccine, we assessed the level of spike-specific multifunctional CD4^+^ T cells in peripheral blood mononuclear cells (PBMC) from vaccinated mice. High frequency of CD4^+^IFN^+^IL-2^+^TNF^+^ were seen after CoVac-II delivery, that were absent from mice vaccinated with spike alone or SpK^Alum^ (Fig. 1C). This corresponds with the Th1-polarising effect of AHQ-II when delivered with inactivated SARS-CoV-2 in animal models and humans^11,14^. We also noted a significant correlation between NAb titres and the level of multifunctional CD4^+^ T cells produced in individual mice (Fig. S2). To further dissect vaccine-induced immunity, we examined the cellular make-up of the draining lymph nodes 7 days after vaccination. Both Alum^SpK^ and CoVac-II induced expansion of spike-specific B cells (CD19^+^MHCII^+^; Fig. 1D) and T follicular helper cell (Tfh) cells (CD4^+^CXCR5^+^PD1^+^; Fig. 1E). The total numbers of antigen specific B cells (Fig. 1F), and Tfh cells (Fig. 1G) were significantly increased in CoVac-II-vaccinated mice compared to immunization with Alum^SpK^. Thus, delivery of trimeric spike antigen with AHQ-II results in strong B and T cell anti-SARS-CoV-2 immune response.

The ability of CoVac-II-induced antibodies to neutralise SARS-CoV-2 variants of concern (VOCs) was examined using isolates of ancestral (Wuhan), as well as Alpha and Beta variants^4^. All plasma samples from CoVac-II-immunized mice neutralised both Alpha and Beta VOCs however titres were reduced for Beta compared to the ancestral virus (8-fold reduction), which is a similar reduction as seen for other COVID-19 vaccines (Fig. 2A)^7^. NAb titres using plasma from SpK^Alum^-vaccinated mice were reduced to the limit of detection against Beta (Fig 2B). Cross-species neutralisation of VOCs was apparent after immunization of rabbits (Fig. 2C) or horses (Fig.2D) with CoVac-II. NAb titres were maintained against the Alpha variant in both species compared to ancestral virus, and high titres against Beta in rabbits (4.7-fold reduction compared to ancestral) and horses (2.7-fold). Thus AHQ-II can adjuvant vaccine immunogenicity across multiple animal models, adding to its already proven immunogenicity in humans as part of the Covaxin vaccine^11^.

**Figure 2.**
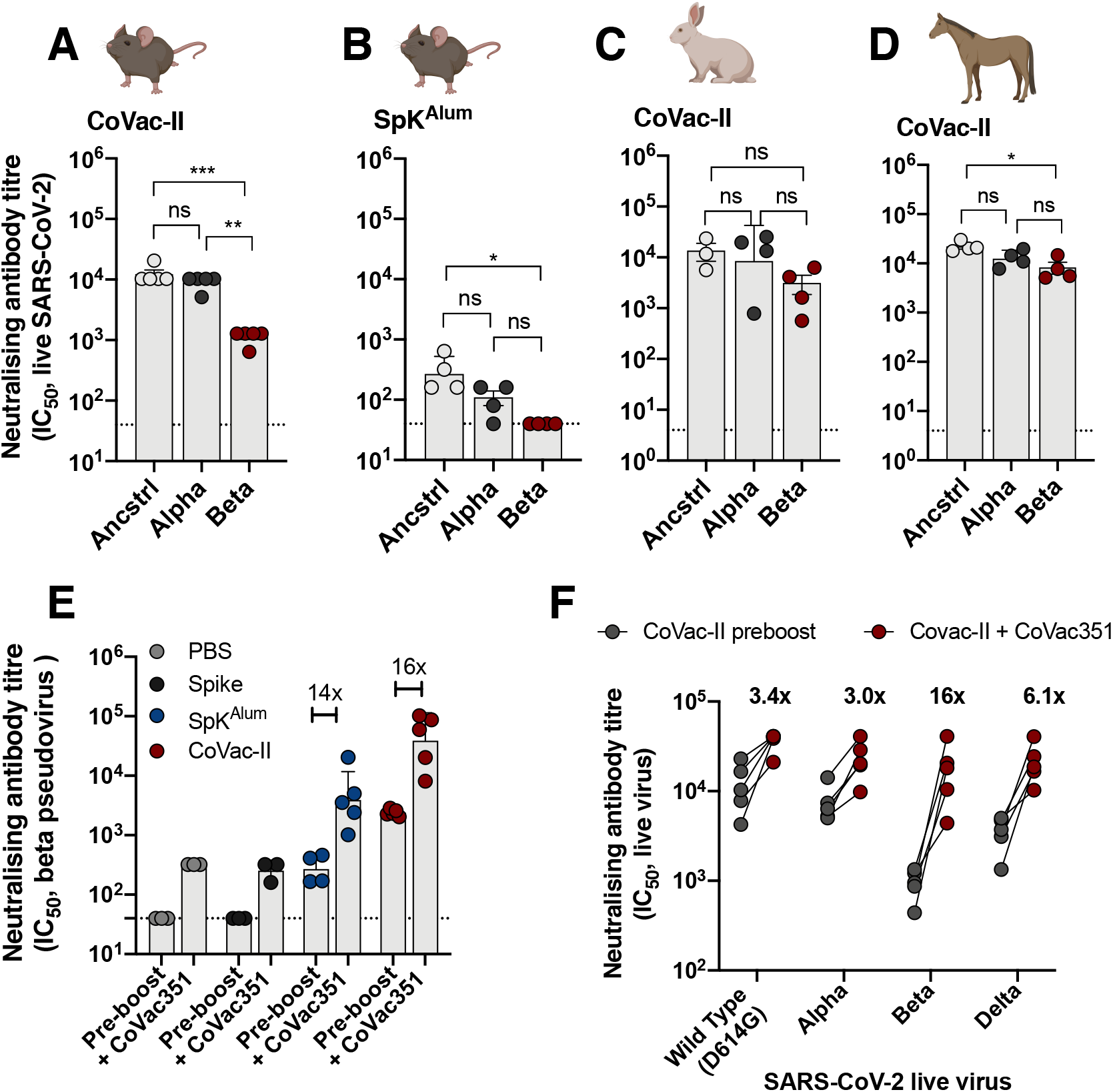
Alhydroxiquim-II-adjuvanted vaccines afford cross-species neutralisation of variants of concern, that is augmented by a variant-specific booster vaccine. Mice (n=4 to 5) were vaccinated as in Figure 1 and 3 weeks post-vaccination, plasma from CoVac-II (**A**) or SpK^Alum^ (**B**) tested for neutralising activity against live SARS-CoV-2 infection of Vero E6 cells (ancstrl=ancestral virus). **C.** Rabbits (n=3) were immunized i.m. twice with CoVac-II (ancestral 5 μg spike/200 μg Alhydrogel) and NAb titres against live SARS-CoV-2 viruses determined. **D.** Horses (n=3) were immunized i.m. twice with CoVac-II (ancestral 20 μg spike/500 μg Alhydrogel) and NAb titres against live SARS-CoV-2 viruses determined. **E.** Mice vaccinated 250 days previously were boosted with a single dose of CoVac351 (5 μg Beta spike/100 μg Alhydroxiquim-II) and NAb titres against Beta spike-pseudovirus determined at one week post-boost. Data presented as geometric mean ± geometric SD. **F.** Plasma from CoVac351-boosted mice was tested for neutralising activity against live SARS-CoV-2 infection of Vero E6 cells. The dotted line shows the limit of detection. Significant differences between groups were determined by one-way ANOVA.

Although CoVac-II immunisation affords some level of cross-neutralisation against the Beta variant, vaccines currently in use display reduced efficacy against this variant when assessed in placebo-controlled or test-negative control trials^8,15,16^. We therefore used our established systems for rapid and high-level production of trimeric viral antigens^10,17^ to produce the Beta spike antigen. Mice vaccinated with CoVac-II eight months earlier (or control groups) were boosted with a single dose of Beta spike formulated in AHQ (CoVac351). Two weeks later, the ability of their plasma to neutralize Beta spike-pseudotyped virus was determined^18^. The increase of NAbs in response to this booster injection was greatest in mice previously vaccinated with SpK^Alum^ or CoVac-II (approximate increase of 16-fold compared to pre-boost levels), however responses were maximal in the CoVac-II prime, CoVac351 boosted group (Fig. 2E). High numbers of spike-specific, multifunctional Th1 CD4^+^ cells were observed in PBMCs from the CoVac-II/CoVac351 group (Fig. S3). To determine breadth of cross neutralisation afforded by boosting with the Beta variant, plasma from CoVac-II-primed, CoVac351-boosted mice was examined for neutralisation of ‘wild-type’ virus and three VOCs; Alpha, Beta and Delta. In pre-boost samples, all VOCs were neutralised, however titres were reduced approximately 10-fold against Beta compared to ‘wild-type’ virus (Fig. 2F). However boosting with CoVac351 resulted in enhanced NAb titres against all VOCs, with greatest increase seen against the Beta variant (Fig 2F). Notably, neutralisation titres against the Delta variant were high (>10^4^) with only a small reduction in titre (approximately 2.2-fold) compared to ‘wild-type’ virus. Of note, these NAb titres are at least 1 order of magnitude higher than the average human convalescent response, which we have previously assessed with similar methodology, and are comparable with that of ‘elite’ neutralisers^4^.

In conclusion, the CoVac-II subunit vaccine we described in this report demonstrates remarkable longevity of immune responses (no decay in NAbs up to 8 months post-vaccination in mice) and is highly immunogenic in multiple animal models, including rabbits and horses. The waning of immunity observed in convalescent patients^4^ and with current vaccines, coupled with low NAb titres correlating with breakthrough infections^19^, suggests that maintenance of humoral immunity will be critical to ensure prolonged vaccine-induced protection against disease. NAbs developed in all immunised species are able to effectively neutralise SARS-CoV-2 variants of concern, which can be augmented by boosting with variant-specific spike vaccines. CoVac-II-immunity compares favourably with other vaccines tested in the same models, that have subsequently shown high-levels of protection in humans^13,20,21^. The excellent safety profile and immunogenicity demonstrated by a AHQ-II-adjuvanted inactivated SARS-CoV-2 vaccine^11^, coupled with our ability to manufacture multiple, high quality antigens at scale ^10^, suggests that AHQ-II/spike protein combinations could constitute safe, affordable and mass-manufacturable COVID-19 vaccines for global distribution.

## METHODS

### Immunization

Female C57BL/6 mice (6-8 weeks of age) were purchased from Australian BioResources (Moss Vale, Australia), and housed at the Centenary Institute in specific pathogen-free conditions. All mouse experiments were performed according to ethical guidelines as set out by the Sydney Local Health District (SLHD) Animal Ethics and Welfare Committee.

SARS-CoV-2 full-length spike stabilized, trimeric proteins were expressed in CHO cells and purified as previously described^10^. Mice (n=3-5) were vaccinated subcutaneously (s.c) once with 5 μg of spike combined with 100 μg of Alhydrogel (Alum; Invivogen, California, USA) or 100 μg Alhydroxiquim-II and boosted three weeks after the first vaccination. In some experiments, mice were further boosted with 5 μg B.1.351/Beta spike protein in 100 μg Alhydroxiquim-II. Mice were bled fortnightly after the first immunization and plasma was collected after centrifugation at 300 × *g* for 10 min. Remaining blood was resuspended in PBS Heparin 20 U/mL, stratified on top of Histopaque 10831 (Sigma-Aldrich, Missouri, USA) and the PBMC layer collected after gradient centrifugation.

Rabbit and horse immunization experiments were performed at Envigo/Cocalico Biologicals (Reamstown, PA, USA), and at East Tennessee Clinical Research, Inc. (Rockwood, TN, USA), respectively, in accordance with institutional guidelines. Adult New Zealand White rabbits (n=4 per group) were bled for pre-immune sera via the marginal vein of the pinna, and immunized intramuscularly (i.m) in the flank region on Days 1, and 15 (Prime + One-boost regimen) with 5 μg of spike protein, adjuvanted with 200 μg (Al content) of either Alhydroxiquim-II, or Alhydrogel, each formulated in 0.2 mL saline. Horses (n=3) were immunized i.m. in the neck region on Days 1, and 15 with 20 μg spike/500 μg Alhydroxiquim-II, or Alhydrogel.

### Flow cytometry assays

To examine T and B cell populations, cells were collected from popliteal lymph nodes 7 post immunization and surface stained (2×10^6^ cells) with Fixable Blue Dead Cell Stain (Life Technologies), spike-AF647 (1 μg) and/or the antibodies listed in Supplementary Table 1. Where required, Cells were then fixed and permeabilized using fixation/permeabilization kit (ThermoFischer) according to the manufacturer’s protocol. To assess spike-specific cytokine induction by T cells, murine PBMCs were stimulated for 4 hrs with spike (5 μg/mL) and then supplemented with Protein Transport Inhibitor cocktail (Life Technologies, California, USA) for a further 10-12 hrs. Cells were surface stained with Fixable Blue Dead Cell Stain (Life Technologies) and marker-specific fluorochrome-labeled antibodies (Supplementary Table 1). Cells were then fixed and permeabilized using the BD Cytofix/Cytoperm^TM^ kit according to the manufacturer’s protocol and intracellular staining was performed to detect cytokines IFN-γ, IL-2, TNF, IL-17 (see Supplementary Table 1 for details). All samples were acquired on a BD LSR-Fortessa (BD) or a BD-LSRII and assessed using FlowJo^TM^ analysis software v10.6 (Treestar, USA).

### Live SARS-CoV-2 and pseudovirus neutralisation assay

High-content fluorescence microscopy was used to assess the ability of plasma from mice to inhibit live SARS-CoV-2 infection and the resulting cytopathic effect in live permissive cells (VeroE6, final MOI=0.05), using previously described methodology^4^. Alpha (B.1.1.7), Beta (B1.351) and Delta (B.1617.2) variants were compared against ‘wild-type’ D614G virus from the same clade (B.1.319), or ancestral virus (Wuhan). The cut-off for determining the neutralisation endpoint titre of diluted serum samples was set to ≥50% neutralisation. Neutralising antibody titers in rabbit and horse immune sera were quantified at Virovax using an automated, liquid-handler-assisted, high-throughput, microfocus neutralisation/high-content imaging methods developed at ViroVax. Briefly, serially-diluted sera (paired pre-immune and immune) and SARS-CoV-2 virus (final MOI of 10) were added to 10^6^/mL Vero cells (ATCC® CCL-81) containing 50 μg/mL of propidium iodide (PI) and plates were then loaded in an IncuCyte S3 high-content imaging system (Essen Bioscience/Sartorius, Ann Arbor, MI). Longitudinal image acquisition and processing for virus-induced cytopathic effect (CPE) and cell death (PI uptake) were performed every six hours, until cell death profiles had crested and stabilized (3.5 days). Neutralising antibody titers (expressed as IC_50_) were obtained from four-parameter logistic curve-fits of cell death profiles using OriginPro 9 (Origin Lab Corp., Northampton, MA).

Replication-deficient SARS-CoV-2 Spike pseudotyped lentivirus particles were generated by co-transfecting GFP-luciferase vector and ancestral^4^ or Beta^18^ spike expression constructs with lentivirus packaging and helper plasmids into 293T cells using Fugene HD (Promega) as previously described ^22^. To determine neutralising antibody titres, pseudovirus particles were incubated with serially diluted plasma samples at 37°C for 1 hr prior to spinoculation (800xg) of ACE2 over-expressing 293T cells^22^. 72 hr post-transduction, cells were fixed and stained with Hoechst 33342 (NucBlue™ Live ReadyProbes™ Reagent, Invitrogen) as per the manufacturers instruction’s, imaged used an Opera Phenix high content screening system (Perkin Elmer) and the percentage of GFP positive cells was enumerated (Harmony® high-content analysis software, Perkin Elmer).

### Statistical analysis

The significance of differences between experimental groups was evaluated by one-way analysis of variance (ANOVA), with pairwise comparison of multi-grouped data sets achieved using Dunnett’s *post-hoc* test. Differences were considered statistically significant when p ≤ 0.05.

### Data availability

The results supporting the findings in this study are available upon request from the corresponding authors.

## Acknowledgements

This work was supported by MRFF COVID-19 Vaccine Candidate Research Grant 2007221 (CC, MS, SGT, JAT). ViroVax LLC gratefully acknowledges funding by the National Institutes of Health (NIAID Contracts HHSN272201400056C (Adjuvant Discovery Program) and HHSN272201800049C (Adjuvant Discovery Program), enabling the discovery and development of the Alhydroxiquim-II, as well as supplemental funding (HHSN272201800049C) for SARS-CoV-2 work. The following reagents were deposited by the Centers for Disease Control and Prevention and obtained (ViroVax) through BEI Resources, NIAID, NIH: SARS-Related Coronavirus 2, Isolate USA-WA1/2020, NR-52281; hCoV-19/England/204820464/2020, NR-54011; hCoV-19/South Africa/KRISP-EC-K005321/2020, NR-54008; hCoV-19/South Africa/KRISP-K005325/2020; NR-54009; hCoV-19/Japan/TY7-503/2021 (Brazil P.1), NR-54982. We thank Charles Baily, Centenary Institute, Sydney for provision of lentivirus packaging and helper plasmids. We acknowledge the support of the University of Sydney Advanced Cytometry Facility and the animal facility at the Centenary Institute. Images created with Biorender.com.

## Contributions

CC, MJW, SGT, MS, FMW, SAD and JAT designed the study. CC, PP, AOS, CA, HL, NDB, AA, MS, SD performed the experiments. All authors contributed to data analysis/interpretation. TT, BMD, NRL provided support in methodology and resources for this study; JAT wrote the first manuscript draft and all authors provided revision to the scientific content of the final manuscript.

---

**Supplementary Figure 1.**
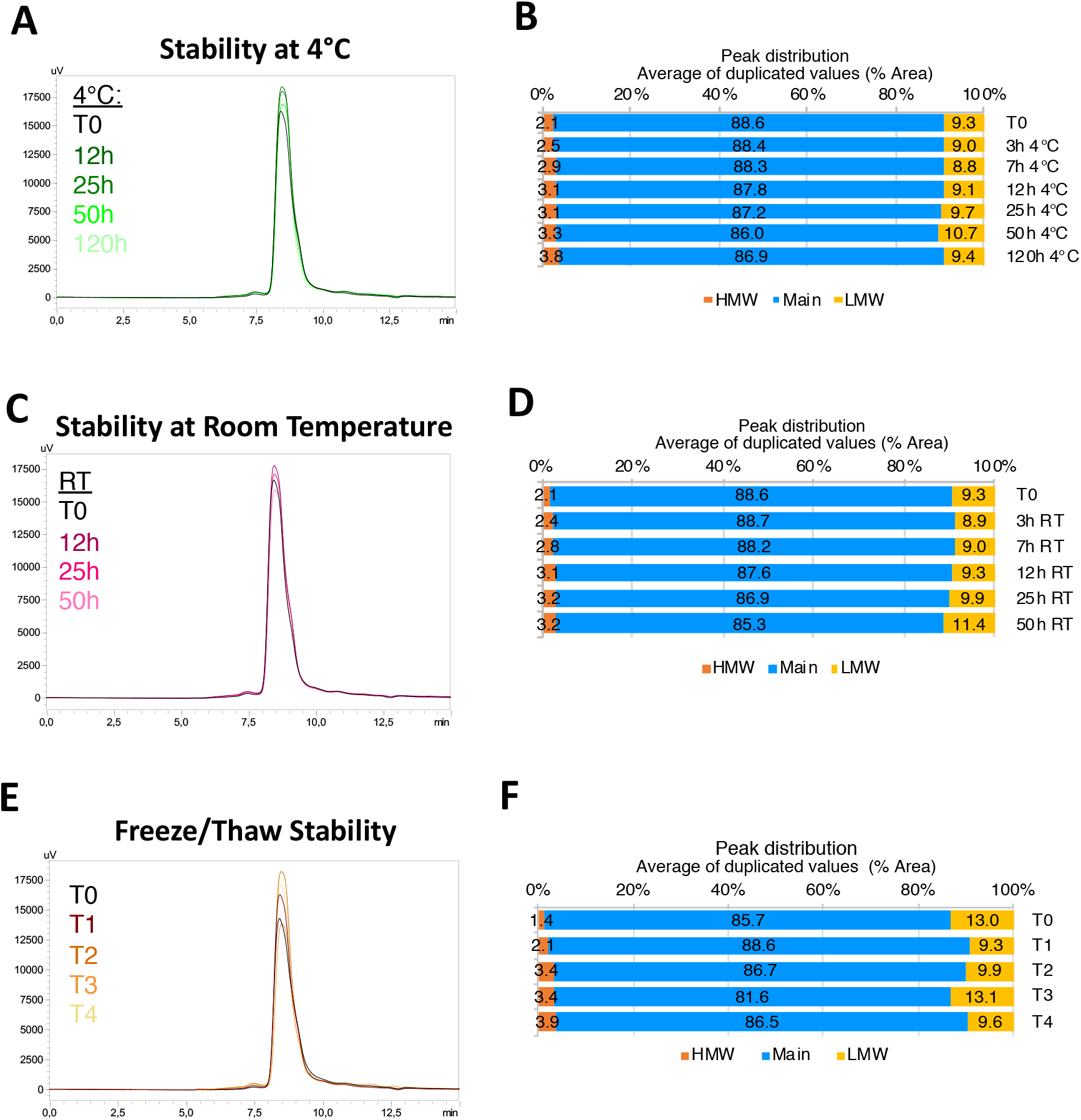
Stability of trimeric spike protein produced in CHO cells. Chromatographs (**A, C, E**) and peak distribution (**B, D, F**) of spike protein after storage at 4°C (**A, B**) and room temperature (RT, **C, D**) or after 4 cycles of freeze-thawing (**E, F**).

**Supplementary Figure 2.**
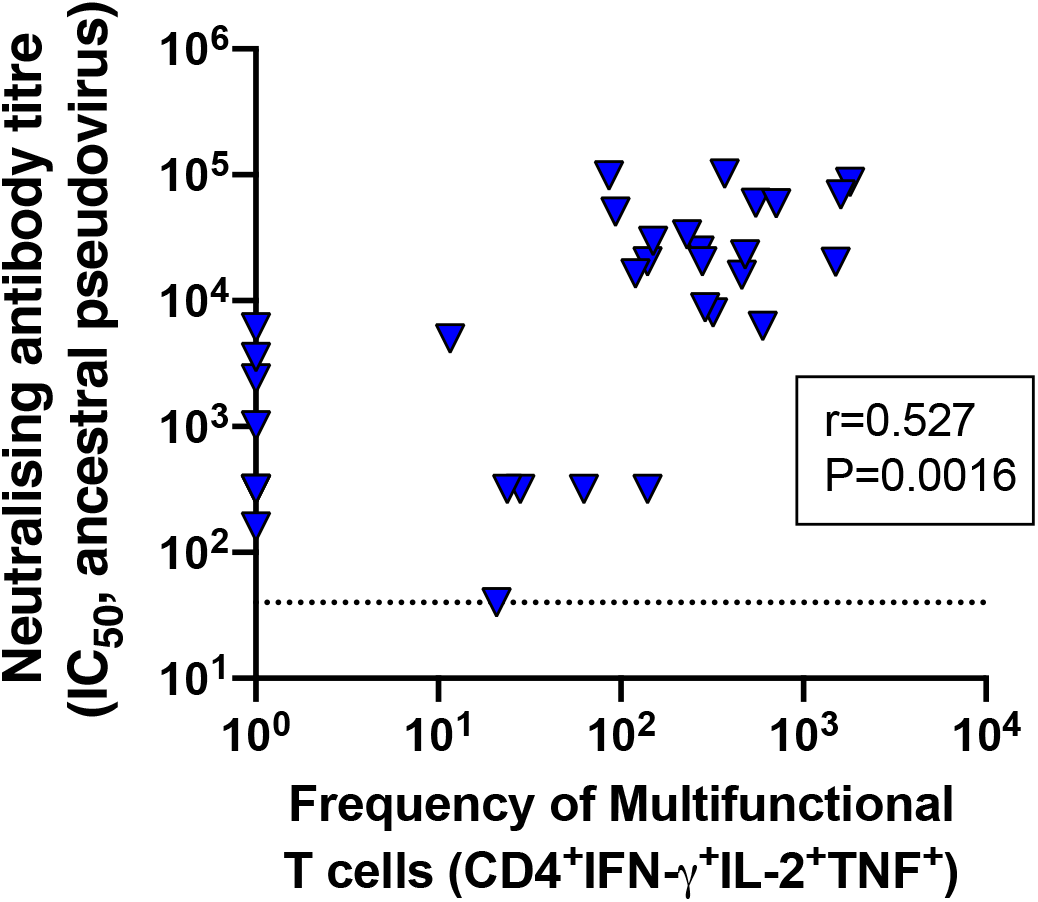
Spearman correlation of neutralizing antibody titre and multifunctional CD4^+^ T cells after vaccination with AHQ-adjuvanted spike vaccines. The dotted line shows the limit of detection.

**Supplementary Figure 3.**
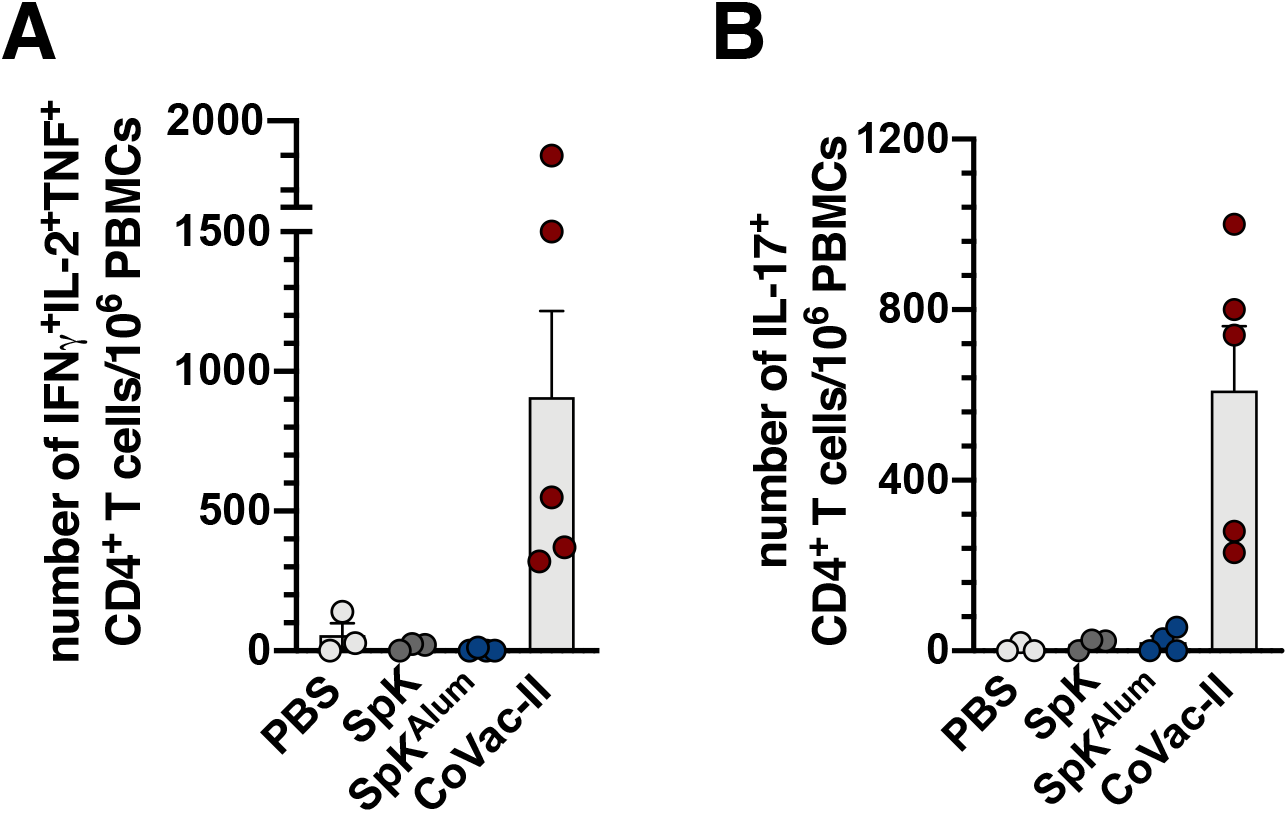
Mice vaccinated 250 days previously were boosted with a single dose of CoVac351 (5 μg Beta spike/100 μg Alhydroxiquim-II). PBMCs isolated one week post-boost were restimulated *ex vivo* with 5 μg/mL of Beta spike and the number of circulating CD4^+^ expressing IFN-γ, IL-2 and TNF (**A**) or IL-17 (**B**) determined by flow cytometry.

